# Decoding Bispecific Antibody Developability: Design Rules and Predictive Models from a 160-Member Library

**DOI:** 10.64898/2026.06.15.732449

**Authors:** Seth Ritter, Laura Rand, Sai Karthick, Tim Bloomingdale, Alexander Smith, Xiang Ao, Yolaine Pierre, Blake Harris, Joshua Moller, Aanal Bhatt, Rebecca Bhatt, Jacob Schwartz, Lucia Grippo, Rich Cohen, David W. Borhani, Peter M. Tessier, Ammar Arsiwala

## Abstract

Bispecific antibodies deliver functional outcomes that monospecific antibodies cannot, yet emergent self-association, polyreactivity, and aggregation often degrade their developability relative to their parental arms. Whether bispecific developability inherits from the parents or is driven by the format has not been tested at scale. We characterized 160 bispecific antibodies and their 65 parental arms on a uniform knobs-into-holes CrossMab IgG1 scaffold across 10 assays on the PROPHET-Ab high-throughput platform. Bispecific developability separates into three classes of inheritance. Hydrophobicity and surface charge inherit cleanly from the parents (Spearman ρ ≈ 0.85 to 0.95), so parental-level screening predicts bispecific fate.

Self-association and polyreactivity inherit partially (ρ ≈ 0.60 to 0.88), with mechanistically interpretable emergent outliers driven in part by Fv-Fv charge complementarity and a parental biophysical ceiling on the hydrophobicity (HIC) by surface-charge (HAC) plane. Thermostability is poorly predicted from parental antibodies (ρ < 0.4), so it requires bispecific-level testing. The class framework yields actionable selection rules: triage hydrophobicity and charge at the parental level, avoid pairing two high-HIC x high-HAC arms, pair opposite-sign Fv charges to suppress self-association but re-validate at the formulation buffer, and measure thermostability on the bispecific itself. This work charts a tractable path from monospecific sequence to bispecific developability prediction.

**Significance:** Bispecific antibodies are a fast-growing therapeutic class, yet the rational design of well-behaving bispecific antibodies from validated monospecific antibody building blocks remains challenging. A key bottleneck is the lack of comprehensive, high-quality public datasets linking parental antibody developability properties to corresponding bispecific antibody developability properties. We address this gap by releasing a dataset comprising 160 bispecific antibodies and the 65 parental monospecific antibodies profiled in 10 developability assays. The data show that bispecific antibody developability is complex. Some properties are easily predictable from the parents, whereas others emerge in the bispecific format or from the bispecific format itself. The factors that govern each property can be identified empirically and used to make practical selection decisions. The mechanistic explanations and predictive models reported here establish a compact set of actionable rules. Together, they define a framework for using computational pipelines to convert monospecific antibodies into bispecific antibodies with drug-like developability properties, enabling faster and more effective generation of high-quality bispecific antibodies for diverse therapeutic applications.

## Introduction

Bispecific antibodies bind two distinct targets simultaneously. This dual engagement enables therapeutic mechanisms unavailable to monospecific antibodies alone, such as modulating two signaling pathways at once, redirecting immune effector cells to tumors, and bridging cell-surface receptors to drive proximity-induced signaling (Labrijn et al., 2019; Klein et al., 2024). Fourteen bispecifics are approved in oncology, immunology, and ophthalmology. The clinical promise of bispecifics is evident from development pipelines: As of 2026, more than 200 bispecific candidates are in clinical development with the number of programs growing 25% annually (Klein et al., 2024). Bispecifics have moved from being a niche engineering curiosity to a mainstream therapeutic modality.

The practical reality, however, is that bispecific development often falls short of the additive promise. Monospecific antibodies with favorable developability profiles (Jain et al., 2017; Bailly et al., 2020) routinely combine to form bispecifics that exhibit emergent properties not predicted by the parents, including increased self-association, polyreactivity, and aggregation, as well as compromised thermal stability (Brinkmann & Kontermann, 2017; Spiess et al., 2015). The mechanistic origin of these emergent properties has been described primarily for individual cases (Brinkmann & Kontermann, 2017). Engineering solutions to pairing two parental antigen-binding fragments (Fabs, herein referred to as “arms”) have substantially improved the success rate of bispecific assembly: for heavy chains, knobs-into-holes (KIH) heterodimerization (Ridgway et al., 1996) and charge-pair steering (Gunasekaran et al., 2010; Lewis et al., 2014); and common light chains (Merchant et al., 1998) and CrossMab pairing (Schaefer et al., 2011). A key question remains unanswered, however: Which developability properties of a bispecific antibody can be predicted from those of its parents, and which must be measured directly?

Resolving that question is complicated by two issues. First, the developability profile of a bispecific antibody could be an additive effect of the constituent parts (Fabs, or variable regions [Fvs]), an inherent effect of a particular bispecific scaffold, an emergent effect of interactions that occur only in the bispecific, or some combination of all three. Second, the data needed to begin disentangling these possible effects are lacking: The combinatorial space of bispecifics implied by the ∼900 approved or clinical-stage monospecific antibodies (Crescioli et al., 2025) approaches millions, yet only individual case studies rather than systematic investigations linking parental developability to bispecific outcomes have been published.

Here we provide the first such comprehensive dataset that compares the developability properties of parental monospecific antibodies to the corresponding filial bispecific antibodies, using our automated developability platform, PROPHET-Ab, which generates standardized, machine-learning-ready data across more than a dozen biophysical assays at a throughput of thousands of antibodies per week (Arsiwala et al., 2025). We studied a deliberately diverse 160-member bispecific antibody library derived from 65 unique parental antibodies already characterized using PROPHET-Ab, built on a standard 1x1 bispecific IgG1-like scaffold. Our dataset allows us to begin to answer the key question—is it the parents, or the child?—and sets the stage for the development of approaches, heuristics, and improved computational tools that will enable the more reliable creation of well-behaving bispecific antibodies.

## Results

### Assaying the developability of 160 bispecific antibodies and their 65 parents

We profiled 160 bispecific and the corresponding 65 monospecific parental antibodies on the PROPHET-Ab developability platform (Arsiwala et al., 2025) (**Figure 1**). The subset of parental antibodies that maintained IgG1 isotype (as used in this study) were compared with measurements from GDPa1, finding high platform reproducibility (Pearson r 0.86-1.00, **Figure S1**). Bispecific antibodies were constructed using an engineered, asymmetric IgG1 scaffold that combines C_H_1 and a parental Fv + C_L_ (kappa or lambda) to form each Fab, a CrossMab C_H_1-C_L_ chain crossover, and Fc modifications to foster proper assembly (see Methods for details).Only one of the two possible arrangements was chosen for each pair of parental antibodies. Parental antibodies were constructed using a standard IgG1 scaffold, regardless of the original isotype, to eliminate confounding isotype-driven differences.

**Figure 1.**
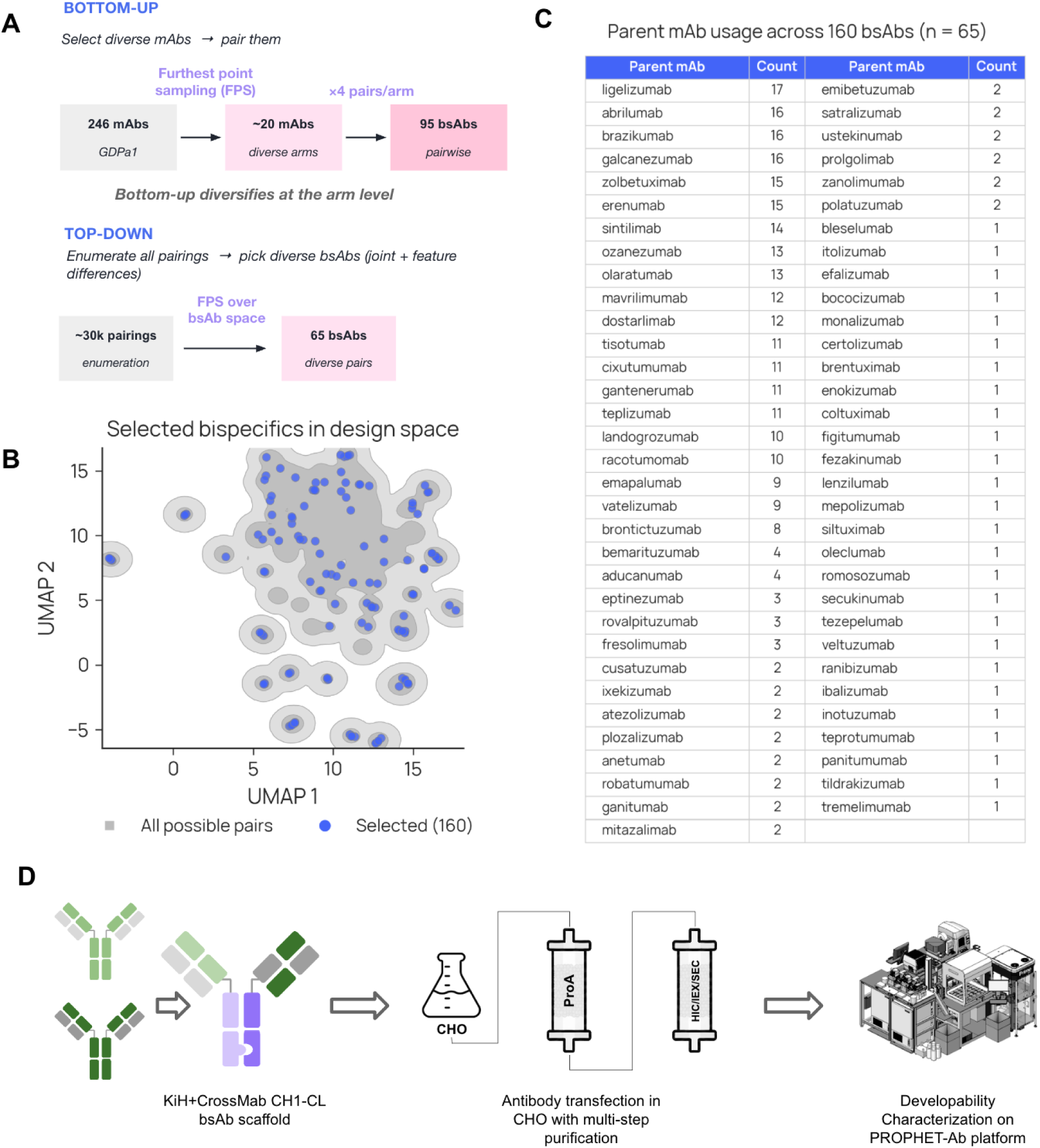
Bispecific library design and experimental workflow. **(A)** Outline of the two sampling strategies used to combine parental antibodies into bispecifics. **(B)** UMAP projection of the bispecific design space using the computed and measured features. Gray: all possible pairings from the GDPa1 panel. Blue: the 160 selected bispecifics. **(C)** Number of times each monospecific parent appears across the 160-bispecific library. The bottom-up strategy reuses each selected parent across multiple partner contexts; the top-down strategy broadens parent coverage to 65 total. **(D)** Experimental pipeline for profiling bispecific antibody developability. Bispecifics were assembled on a multi-engineered KiH/CrossMab IgG1 scaffold, expressed via transient transfection in CHO cells, purified by Protein A capture followed by polishing steps, and profiled across 12 assays (10 developability, 2 quality) on the PROPHET-Ab platform.

The 160 bispecifics represent approximately 8% of the 2,080 possible ways to pair 65 parental antibodies, which were selected from the 246-antibody set we previously characterized (Arsiwala et al., 2025). We used a deliberate pair selection strategy in order to sample a diversity of outcomes. Two complementary farthest-point sampling approaches spanned a combined experimental property and *in silico* feature space: one maximized the diversity of the parents, and the other the diversity of the resulting bispecifics. This selection strategy maximized the biophysical diversity we were able to sample within a fixed experimental budget, but it precluded evaluation of population-level statistics of arbitrary (30,135 total) pairings of the 246 antibodies (**Figure 1, A-C**).

All antibodies were produced via transient expression in Chinese hamster ovary (CHO) cells, followed by Protein A affinity capture and up to three polishing purification steps (**Figure 1, D**). Production quality-control metrics (yield, concentration, endotoxin, and purity) were recorded for each sample.

Antibodies were profiled in 10 developability assays spanning self-association, surface properties, polyreactivity, and thermostability (Arsiwala et al., 2025). The assays comprised: Affinity-capture self-interaction nanoparticle spectroscopy (AC-SINS), which measures self-association propensity, performed in three conditions (PBS, pH 7.4, histidine/arginine, pH 6, and histidine/NaCl, pH 6; three chromatographic assays which measure surface properties, heparin affinity chromatography (HAC), hydrophobic interaction chromatography (HIC), and stand-up monolayer adsorption chromatography (SMAC); polyreactivity, which measures nonspecific binding, to CHO cell lysate, ovalbumin, and baculovirus particles (BVP); and nanoscale differential scanning fluorimetry (nanoDSF) which measures thermal unfolding transitions which we labeled as Tm1, Tm2. Additionally, quality-control and characterization readouts included purity measured by size-exclusion chromatography (SEC, reports percent monomer) and intact mass spectrometry (IntactMS, reports bispecific purity).

### The asymmetric bispecific scaffold does not substantively impact most developability properties

To evaluate the impact of the asymmetric bispecific scaffold on developability properties, we produced 21 bispecific antibodies in both possible orientations and compared pairwise the properties of these 42 antibodies (**Figure 2**). In adsorptive chromatography (HIC, SMAC, HAC), self-association (AC-SINS, three conditions), polyreactivity (PR-CHO, PR-Ovalbumin, PR-BVP) assays, the bispecific antibody pairs behave similarly regardless of orientation (Spearman correlation coefficients, ⍴, ranging from 0.78 to 0.95; n = 21). Thermostability (Tm1, Tm2), by contrast, correlates weakly, possibly due to differing inter-domain interactions inherent to the scaffold. These data provide evidence that most biophysical developability properties are invariant to the asymmetry of the chosen bispecific orientation.

**Figure 2.**
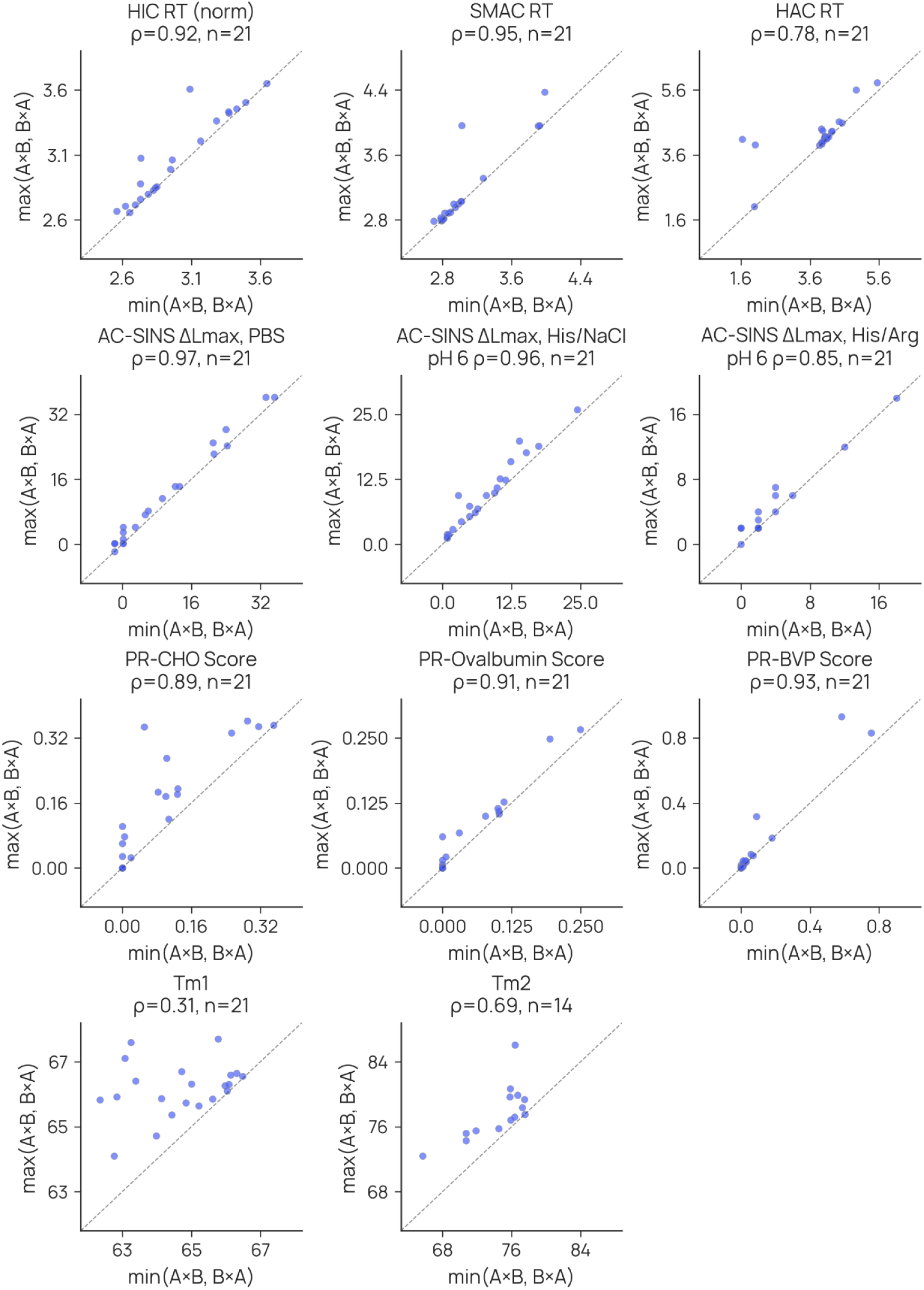
Orientation swap analysis of parents. Measurements of bispecifics where both orientations are present (e.g. AxB and BxA) are shown. For permutation consistency, x-axis is the minimum and y-axis is the maximum. Most assay readouts, excluding thermostability, exhibit high correlations with few outliers.

### Bispecific antibodies predictably inherit most developability properties from their parents

A bispecific antibody joins two different Fvs into a single molecule. If developability properties are set largely by those of the constituent Fvs, some simple combination of the property values of the two parental antibodies should predict the corresponding value for the bispecific antibody. Three combining rules—mean, min, and max—yield predictive models with the least prior assumptions: both parents contribute equally, or one parent dominates. We tested this notion across the developability panel: for each assay, we correlated the property values observed for bispecific antibodies with those predicted by combining the parental antibody property values. For each assay (n = 125 to 160), we report the Spearman ρ of the three combining operators (**Figure 3**).

**Figure 3.**
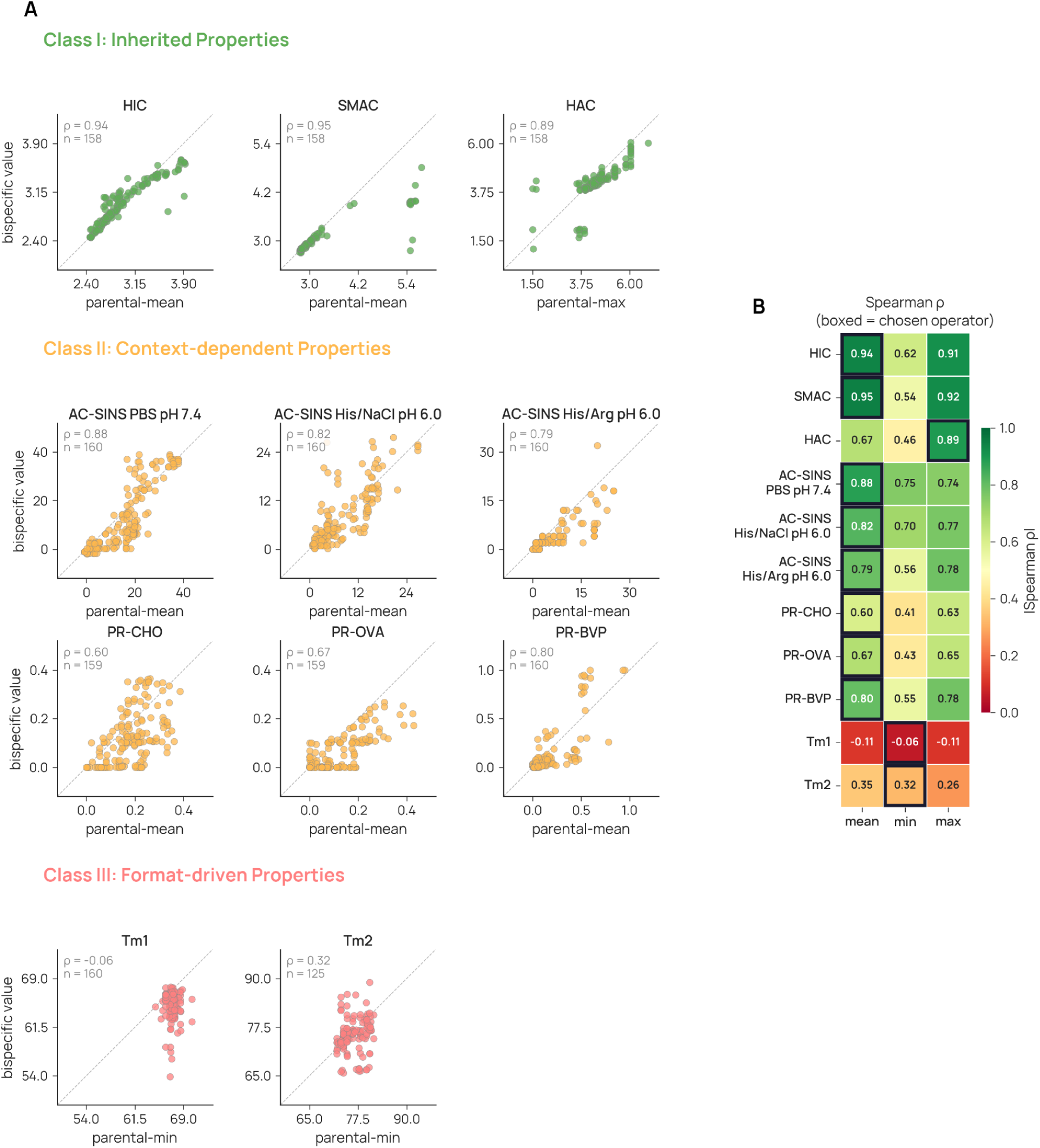
Bispecific antibodies predictably inherit most developability properties from their parents. **(A)** Scatterplots of operator-predicted versus observed bispecific values for representative assays, organized into three classes. **(B)** Heatmap of Spearman’s ρ values between parental combinations and observed bispecific values for three operators (mean, min, max) across all assays. Rows are sorted by the best operator’s ρ. Highlighted cells indicate the chosen operator per assay. n = 125-160 depending on assay coverage.

We found that the predictability of bispecific antibody developability properties from those of the parents varies continuously across the 10 assays, with ρ ranging from 0.95 to –0.11. For 9 of 11 assay readouts, the mean of the parental property values best predicted the bispecific value, but the maximum performed nearly as well; the minimum gave poorer predictions. For thermostability, however, we used the minimum, which performs similarly to the mean, as it is more consistent with the underlying physical mechanism, i.e., relatively independent unfolding of domains.

The developability properties a bispecific antibody most predictably inherits from its parents are the measures of surface hydrophobicity and (positive) charge, HIC, SMAC, and HAC (ρ 0.95, 0.94, 0.89). HAC is the one property for which the maximum provides a (much) better prediction than mean (ρ 0.89 vs. 0.67). Self-association (AC-SINS) is nearly as well-inherited, with some measurement conditions yielding somewhat better predictions (PBS pH 7.4, ρ 0.88) than others (His/Arg pH 6.0, ρ 0.79). Polyreactivity is also reasonably predictable for PR-BVP (ρ 0.80), but the other two proved less so (ρ 0.67, 0.60); the reasons for this difference are not clear. Finally, the thermostability of a bispecific antibody is very poorly predicted from those of its parents (ρ < 0.4), in line with our observations in the arm-swap control experiment. We discuss these three classes of inheritance below.

### Class I: Bispecific antibody properties that are easily predicted from the parents

Bispecific antibody surface properties—hydrophobicity and (positive) charge—as measured by the adsorptive chromatography methods HIC, SMAC, and HAC, are well-predicted (ρ ≥ 0.89) from the properties of the parental antibodies (**Figure 3**).

Use of the mean combining operator posits that the contributions of each Fv to the overall surface properties of the bispecific antibody are additive. For measures of hydrophobicity (HIC and SMAC), this idea seems intuitively reasonable, and is borne out in practice. The maximum performs nearly as well, however, suggesting that interaction with the chromatographic matrices is driven by the most-hydrophobic portion of the antibody

HAC measures binding to immobilized heparin, a negatively charged carbohydrate that mimics the negatively charged surfaces of cell membranes (Kraft et al., 2020). Somewhat surprisingly, the maximum combining operator gave better predictions (ρ 0.89) than did the mean (ρ 0.67). This result suggests that positively charged surface patches on the Fv which binds heparin more strongly dominate the association of a bispecific antibody with heparin, rather than both Fv contributing proportionately.

These results establish a parent-level screen for adsorptive chromatographic properties. Because bispecific antibody HIC, SMAC, and HAC values are easily predicted (ρ ≥ 0.89) from parent measurements, these properties can be triaged at the monospecific stage without assembling the bispecific format.

### Class II: Bispecific antibody self-association is inherited but is pairing-sensitive and buffer-dependent

Self-association is also inherited, but for two subsets of antibodies it is sensitive to the pairing context. Across the three AC-SINS buffer conditions, the mean of the two parental values predicted bispecific self-association propensities (Spearman ρ 0.79–0.88), revealing that most bispecifics track their parents. As shown in **Figure 4A**, however, two subsets deviated by more than expected from replicate variation alone.

**Figure 4.**
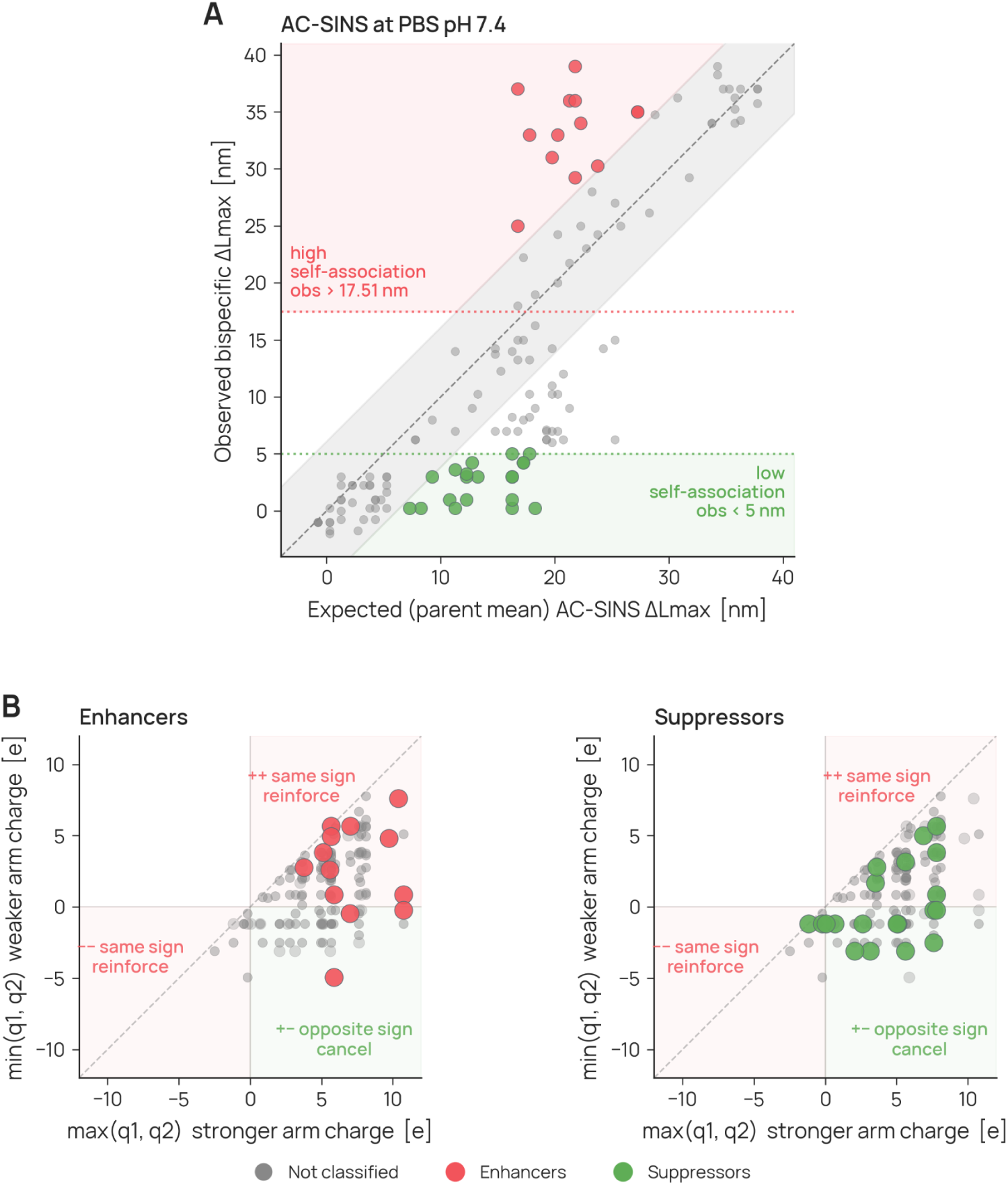
AC-SINS self-association in PBS pH 7.4. Enhancer and suppressor mechanism. **(A)** Observed vs parental mean AC-SINS assay value with the noise band and clinical warning threshold (red dotted line, derived from Arsiwala 2025) that define enhancers, or below 5 nm (chosen arbitrarily to designate low-self-association) for suppressors. **(B)** Enhancers and suppressors resolved by the two parental Fv’s net charge.

To explore these deviations without implying a mechanism, we classified bispecific pairings as self association-enhancing (red; undesirable) or self association-suppressing (green; desirable) relative to a noise envelope, around the idealized inheritance, derived from replicate standard deviations of both parental and bispecific antibodies. Enhancing pairings exhibit self-association higher than anticipated from parentals values (**Figure 4A**); suppressing pairing exhibit opposite behavior. In the PBS pH 7.4 condition, for example, this classification identified 13 enhancing and 20 suppressing pairings among 139 unique pairs of parental antibodies.

### Charge imbalance drives enhancers, charge balance drives suppressors

We hypothesized that differing Fv electrostatic properties distinguish enhancers and suppressors from bispecific antibodies that display the expected average behavior (**Figure 4B**): for enhancers, same-sign charges reinforce one another to form a large net monopole that drives intermolecular association; for suppressors, opposite-sign charges cancel one another to effectively sequester the antibody from intermolecular association.

To evaluate this hypothesis, we tested the distributional shift of the two outlier groups relative to non-outliers. Six transformations that combine Fv charges in different ways were evaluated. The distributions for the most performant transforms for enhancers compared to average bispecific antibodies, as well as that for suppressors, are presented in **Figure S2**. We found the absolute sum of charge to be the most performant for enhancers (Mann-Whitney-U p = 0.01), and the signed geometric mean to be the most predictive for suppressors (p = 0.02). Together, these results support charge character as one mechanistic contributor to bispecific self-association behavior.

### Suppressors convert into enhancers in different buffer conditions

In some cases, undesirable enhancer behavior emerges only in certain buffer conditions, for example in bispecific antibodies created from brazikumab and gantenerumab, abrilumab and landogrozumab, and ligelizumab and romosozumab. All three are suppressors in PBS pH 7.4 (brazikumab/gantenerumab a super suppressor), but all become strong enhancers in His/NaCl pH 6 (**Figure S3**).

### Class II: Bispecific polyreactivity is pairing-sensitive and amplified by high-HIC × high-HAC arms

We examined the relationship between polyreactivity and surface hydrophobicity (HIC) or positive charge (HAC), as these properties have been suggested to be linked (Raybould et al., 2019). For both parental and bispecific antibodies, those exhibiting high HAC scores tend to also be more polyreactive, independent of reagent, whereas the impact of high HIC is much less pronounced (**Figure S4**). As a visual guide, we highlight a pareto frontier on the HIC x HAC plane (red dotted line) defined by the parental antibodies. Interestingly, several bispecifics cross the HIC x HAC frontier, with combined values beyond any parental antibodies, suggesting “emergence”.

As an extension of this observation on the emergent extremes, we explored the utility of the frontier boundary in HIC x HAC space (**Figure S5A**). We find that bispecifics created from mAbs along this frontier are enriched for polyreactive responses (**Figure S5C**). The distance to this boundary when used as a classifier of the top-20% achieves AUC values between 0.69 and 0.80 depending on the polyreactivity assay (**Figure S5B**).

Individual parental arms illustrate how the hydrophobic axis contributes to this liability. Brazikumab and ligelizumab are high-HIC parental arms, each above the 97th percentile for HIC retention among the monospecific antibodies. However, both have low HAC retention, indicating limited charged-surface character on their own. Thus, these arms contribute primarily to the hydrophobic component of the high-HIC × high-HAC risk state. Bispecifics enter this region when a hydrophobic arm of this kind is paired with a high-charge partner, causing the two parental surface liabilities to combine in the assembled molecule.

### Class III: Bispecific antibody thermostability is format-driven and not predictable

The thermal transition temperatures of bispecific antibodies exhibit poor predictability, with ρ ranging from –0.06 (Tm1) to 0.32 (Tm2) using the minimum combining function (**Figure 3**). Our choice of combining function was guided by the expected mechanism of thermal unfolding. Because individual domains are expected, to a first approximation, to unfold independently, the minimum parental value should best capture the earliest onset of instability in the bispecific antibody. A plausible explanation for the poor predictability of thermostability is that our chosen bispecific scaffold changes the thermal landscape: the asymmetric, domain-swapped Fab and heterodimeric C_H_3 interface may shift, broaden, or merge unfolding transitions relative to the parental antibodies, weakening the predictive relationship. This explanation is supported by our observation that thermostability, especially Tm1, was the developability property most impacted by our arm-swap control experiment (**Figure 2**).

### Complex supervised models do not improve predictability

To determine whether supervised learning could improve upon the baseline provided by the simple (minimum, mean, maximum) combining functions, we evaluated seven regression models across 11 feature configurations and 17 developability labels. Feature sets ranged from simple mean-of-parents representations to multi-assay experimental, in silico, and hybrid configurations. All models were assessed using parent-disjoint leave-one-out cross-validation, ensuring that no test bispecific shared a parent with any training example (**Figure 5A**).

**Figure 5.**
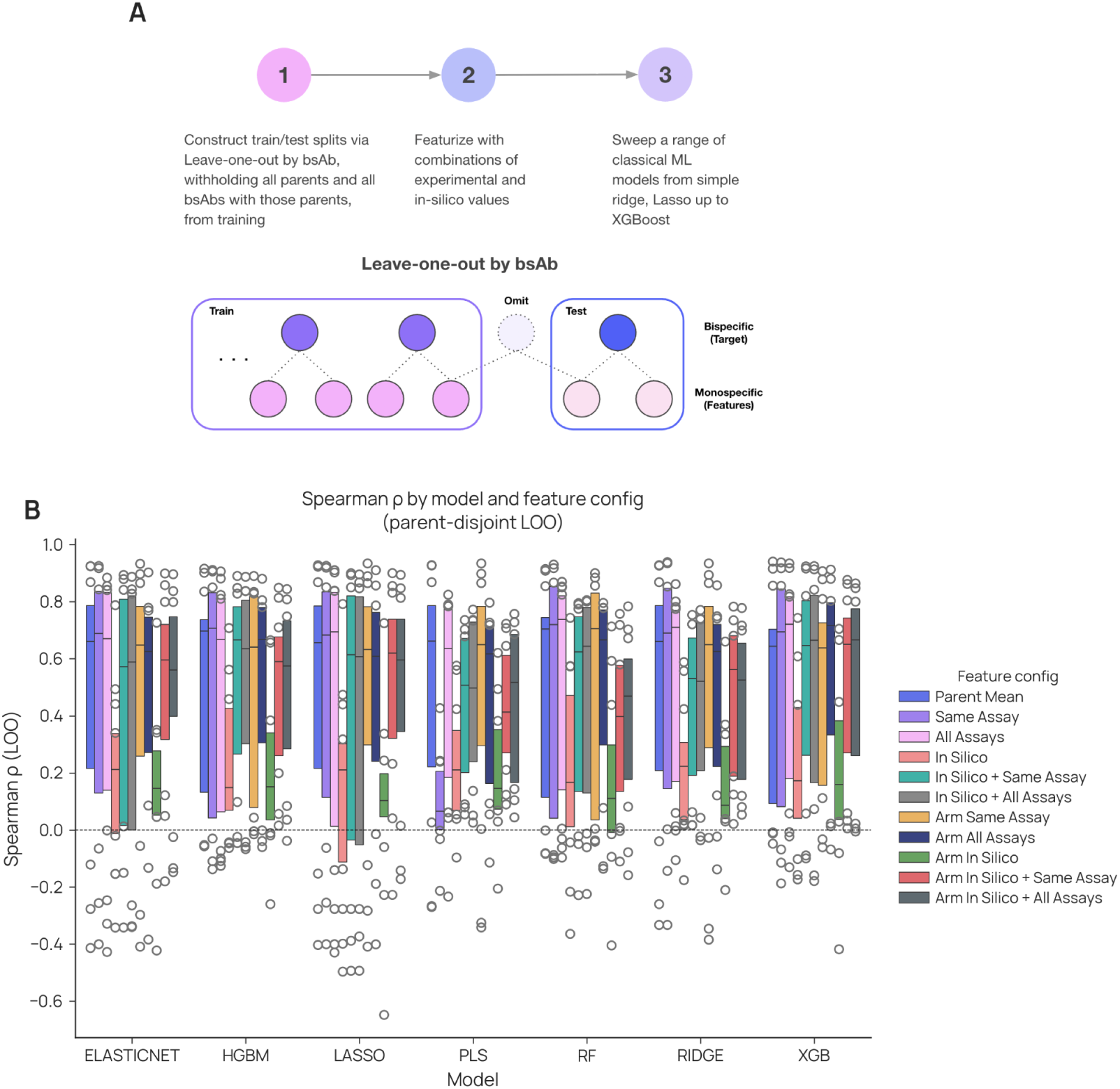
Supervised models do not improve over simple baselines. **(A)** Workflow: parent-disjoint leave-one-out by bispecific (each held-out bispecific and every bispecific sharing either parent are withheld from training), featurization combining experimental and in silico parent values, and a sweep of seven classical regressors from Ridge and Lasso to XGBoost. **(B)** Held-out Spearman ρ by model family and feature configuration across labels; bars are medians, points are per-assay values.

We found that none of these more complex models substantively improved the ability to predict the developability properties of bispecific antibodies. For example, the best-performing models achieved ρ = 0.94 for SMAC, 0.94 for HIC, and 0.87 for HAC (**Figure 5B**), i.e. no improvement over the simple combining functions (0.95, 0.94, and 0.89, respectively). Similarly, the complex models gave no improvement for AC-SINS ρ = 0.76–0.89 (vs. 0.79–0.88 for mean), and only an arguable minor improvement for polyreactivity ρ = 0.70–0.78 (vs. 0.60–0.80 for mean). Thermostability properties remained poorly predictable, with maximum correlations of ρ = 0.24 and 0.41 for Tm1 and Tm2 (vs. –0.11 and 0.35).

### Repeated parent usage identifies parental antibodies that recurrently perturb bispecific developability

The library design allowed us to ask a final question that cannot be answered from single bispecific examples: are some parental Fvs systematically difficult to deploy in a bispecific format? This question is distinct from asking whether a parental antibody has poor developability as a monospecific molecule. A parental Fv could be problematic because it is already poorly behaved on its own, because the bispecific scaffold exposes a liability that is not apparent in the monospecific format, or because it becomes problematic only when paired with particular partners. The repeated use of selected parental Fvs across multiple bispecific pairings therefore provides a way to distinguish recurrent parent-specific effects from isolated partner-specific failures.

To test this, we analyzed the residual behavior of each bispecific after accounting for the value expected from its two parental antibodies. In this analysis, a positive or negative parent-specific effect means that bispecifics containing that Fv tend to shift in the same direction across multiple partners, relative to the parental expectation (**Figure S6**). Thus, the analysis asks whether an Fv carries a reproducible bispecific-format effect that is not explained simply by averaging the two parental measurements.

This analysis identified a small set of high-leverage parental Fvs with recurrent effects across multiple assay conditions. Ligelizumab and gantenerumab showed significant parent-specific effects in seven assay conditions, teplizumab in five, and brazikumab, mavrilimumab, and olaratumab in four. These results indicate that some parental Fvs repeatedly alter bispecific developability across partner contexts. However, these effects should not be interpreted as universal failure with every partner. Rather, they identify parents that require additional scrutiny because their bispecific products deviate reproducibly from the behavior expected from the two monospecific antibodies.

The direction and assay context of these effects are important. Some parent-specific effects correspond to worsened developability, whereas others correspond to apparent suppression in a specific assay. For example, gantenerumab-containing bispecifics showed lower AC-SINS self-association than expected from the parent mean in PBS and histidine/arginine buffer.

The model also revealed assay-wide systemic effects. Thermostability showed a consistent global decrease indicating that the bispecific format systematically reduces thermal stability relative to the expectation, supporting the prior observations of format-driven differences. HAC retention time exhibited a modest positive global shift, whereas HIC showed no significant bias.

## Discussion

Bispecific antibody developability is not a single phenomenon and should not be modeled as one. By studying the developability properties of 160 bispecific antibodies and their 65 parents, we resolved these properties into three classes, based on the degree and nature of the inheritance of each property from the parental antibodies. Though empirical, this classification provides a useful lens through which to discuss developability properties as well as practical engineering choices.

Class I properties are highly predictable, transferring cleanly from parent to bispecific. Hydrophobicity (HIC, SMAC retention) and surface charge (HAC retention) of the bispecific can be predicted from the parents’ properties with a Spearman ρ = 0.85 to 0.95. The bispecific behaves as a blend of its two parents.

Class II properties exhibit moderate predictability but with some context-dependent behavior. Self-association and polyreactivity can be predicted at ρ = 0.60 to 0.88. AC-SINS exhibits two outlier populations: suppressors of self-association, and enhancers. The net charge of suppressor Fvs is more balanced, whereas that of enhancers is not. An interesting effect is observed for a few bispecific antibodies: they are suppressors at pH 7.4, but become enhancers at pH 6, an example of context dependency of inheritance. Similarly, polyreactivity predictivity is context-dependent, depending on the reagent used to measure this property (BVP yields the highest predictivity). Together, these observations are consistent with prior work linking antibody surface hydrophobicity and charge-patch features to nonspecific binding and polyreactivity (Raybould et al., 2019), and with reports that electrostatic surface properties can contribute to antibody self-association (Shan et al., 2018).

Class III properties are not predictable and appear to be driven, at least in part, by the particular bispecific antibody scaffold as has also been observed previously (Chennamsetty et al., 2009; Condado-Morales et al., 2024). The thermostability (Tm1, Tm2) of the bispecific correlates poorly with that of the parents (ρ < 0.35). This finding suggests that the scaffold format alters stability, responding idiosyncratically to the particular Fvs and to their asymmetric arrangement in one of two possible configurations (**Figure 2**). The heterodimeric Fc interface, the engineered C_H_3 mutations, and inter-Fv proximity create unfolding transitions absent from the homodimeric parent antibodies (Amash et al., 2024).

The complex supervised modeling provided no meaningful benefit over using the parental assay values directly via the simple combining functions. Given this transference of parental developability to the bispecific format for most assays we posit that an efficient computational route from sequence to bispecific developability is a two-stage cascade. Stage one converts monospecific sequence into predicted arm-level empirical values (HIC, SMAC, HAC, AC-SINS, polyreactivity). Stage two takes those predicted arm-level values together with format-aware features to predict bispecific developability. Building stage one requires comparable monospecific data at scale, now accumulating through multi-party consortia: FAITE (FAITE Consortium, 2025), Lilly TuneLab (Eli Lilly, 2025), and the Ginkgo-Apheris Antibody Developability Consortium (Ginkgo Datapoints and Apheris, 2025).

Our dataset has limitations that frame how its findings should be used. The 160 bispecifics sample 8% of 2,080 possible pairings of the 65 parents, and the sampling was non-random by design, so conclusions apply to the sampled subset rather than to the full pairing population. All molecules used a single multi-engineered scaffold (knobs-into-holes heterodimerization with a stabilizing disulfide, electrostatic Fc steering, Protein A ablation, and CrossMab CH1-CL crossover on IgG1); alternative architectures (DVD-Ig, BiTE, common-light-chain platforms) impose different constraints and may exhibit different inheritance behavior. All parents were grafted onto a standard IgG1 scaffold, which removes isotype as a confounder but does not reflect native isotype contexts. In silico features were computed on monospecific parental sequences; structure-based predictions on the bispecific heterodimer were not attempted. The format-effect decomposition is limited to additive per-Fv terms, because pairwise (epistatic) terms are not identifiable at the current pair-graph density. The parent-disjoint leave-one-out scheme covers all 160 bispecifics but was trained on overlapping sets that share most data across folds.

From the class framework we extract four design rules, proposed as practical decision support for bispecific discovery campaigns:

1. Triage hydrophobicity and charge at the parental level. Class I assays inherit at ρ ≈ 0.90, so parent-level HIC, SMAC, and HAC screening predicts bispecific fate with high fidelity.
2. Avoid pairing two mAbs that are both high-HIC and high-HAC. In this library, pairings of two such arms increased the probability of exhibiting high polyreactivity.
3. Pair opposite-sign Fv charges to suppress self-association in PBS, then re-validate in the formulation buffer. Histidine protonation at pH 6.0 alters the Fv charge map and can flip a pH 7.4 suppressor into an enhancer. PBS-only assessment systematically misses this risk.
4. Measure thermostability at the bispecific level, as it cannot be predicted from the parents

These results extend rather than overturn the existing literature. Case studies of individual programs have documented emergent self-association, charge variants, and viscosity in bispecific candidates (Wang et al., 2024), and per-Fv format-effect analyses on smaller panels have identified variable regions that carry liabilities across partners. This dataset contributes additional scale and diversity via generation of data for a large set of bispecifics as well as their corresponding parental molecules. By characterizing 160 bispecifics built from 65 distinct parental arms on a standard scaffold, we provide further evidence to the degree of property inheritance and provide a new public dataset for further exploration.

## Conclusions and Future Perspectives

The broader case for this work is straightforward. Bispecific antibody development has remained a primarily empirical exercise because the data needed to make it predictive have not existed at the relevant scale. The 160-member dataset, the class framework, the mechanistic explanations for emergence, and the predictive models reported here, together with the design rules they imply, are intended as a starting point for the field rather than a final word. The goal is to move bispecific antibody developability, as a discipline, from characterization to computation.

## Materials and Methods

### Antibody panel design and production

We assembled a panel of 71 IgG antibodies (GDPa4 monoclonal subset, 65 parents and 6 additional controls) and 160 bispecific antibodies (GDPa4 bispecific subset). Each GDPa4 monoclonal antibody’s variable fragment (Fv) was paired with a uniform IgG1 heavy-chain constant region, regardless of the originator molecule’s native isotype (IgG2, IgG4, murine, or other). Additionally, constant-region types for the light change (κ and λ) were preserved and then manipulated (e.g. via CrossMab swap). This design eliminates isotype as a confounder in both the bispecific panel (which uses a multi-engineered KiH/CrossMab IgG1 scaffold, described below) and the cross-platform comparison with the published GDPa1 benchmark (Arsiwala et al., 2025).

The 160 bispecifics were produced on a multi-engineered IgG1 scaffold (**Figure S7**) combining five heterodimerization and quality strategies: knob-into-hole (KiH) mutations (T366W on the knob chain; T366S, L368A, Y407V on the hole chain) for Fc heterodimerization (Ridgway et al., 1996), a stabilizing interchain disulfide (S354C knob, Y349C hole), electrostatic steering mutations (K409A knob, F405K hole) for additional selectivity, Protein A ablating mutations (H435R, Y436F) on one heavy chain to enable homodimer removal during purification, and CrossMab CH1-CL domain crossover on one arm to prevent heavy-light chain mispairing (Schaefer et al., 2011). Each bispecific comprises four chains (HC1, LC1, HC2, LC2), transfected at a 2:3:2:3 plasmid ratio, and purified by Protein A capture followed by up to three polishing steps. The 160 parent pairings were drawn from the 246 monospecifics in the published GDPa1 panel (Arsiwala et al., 2025) using two complementary farthest-point sampling (FPS) strategies, split approximately equally between approaches. Both strategies operated in a 13-dimensional property space comprising four GDPa1 experimental readouts (HIC retention time, second melting temperature, polyreactivity to CHO cell lysate, and AC-SINS delta Lmax at pH 7.4), five sequence-derived biophysical properties (isoelectric point, hydrophobicity, aromaticity, instability index, and variable-region sequence length), and four structural descriptors from the Molecular Operating Environment (MOE; Chemical Computing Group, 2024) (CDR hydrophobic patch area, ensemble charge, dipole moment, and VH-VL interface binding affinity). All properties were min-max scaled before distance computation.

The bottom-up strategy applied FPS in the monospecific property space to select 20 diverse parents, then constructed bispecific pairs from the selected set. Each parent was paired with two others, and pairings were sampled in both orientations (which parent occupies HC1/LC1 vs. HC2/LC2). This strategy yielded 95 bispecific combinations and ensured repeated observation of each selected parent across multiple bispecific contexts. Twenty one of these pairings were produced in both orientations to enable assessment of arm-position effects on developability.

The top-down strategy enumerated all 30,135 possible bispecific pairs from the full GDPa1 panel and applied FPS in a 26-dimensional space defined by the per-pair mean and absolute difference of each of the 13 properties. After excluding pairs already selected by the bottom-up strategy, FPS selected 65 additional bispecifics that maximized coverage of the bispecific property space. This approach prioritizes diversity in bispecific-level property combinations, complementing the bottom-up strategy’s emphasis on per-parent replication.

The union of both strategies yielded 160 bispecifics constructed from 65 unique monospecific parents, sampling approximately 8% of the C(65, 2) = 2,080 possible pairings among those parents. Distributions of properties (computed and experimental) for the parental antibodies and the anticipated values from the mean for the bispecific are shown in **Figure S8**.

Antibodies were transfected in CHO cells followed by clarification and capture using AmMag Protein A magnetic beads (GenScript) following the vendor’s standard protocol. Bispecific antibodies were further polished by a second chromatography step: HiTrap Butyl HP-HIC (Cytiva 28411005) or HiTrap Q HP-IEX (Cytiva 17101403) or HiLoad 26/600 Superdex 200pg 320ml-SEC (Cytiva 28989336) to remove residual homodimers, half-antibodies, and process-related impurities. Some bispecifics underwent additional polishing as required (see datasheet for details). Final purified material was buffer-exchanged into 20 mM histidine, 150 mM NaCl, pH 6.0 storage buffer and stored at −80 °C. Identity and assembly quality were confirmed by reducing and non-reducing SDS-PAGE (≥95% pure under non-reducing conditions) and analytical SEC-HPLC (≥99% monomer); endotoxin was measured by Limulus amebocyte lysate (LAL) assay and ranged from 0.010 to 0.038 EU/mg across the library. Heterodimer assembly purity for each bispecific was independently verified by intact-mass spectrometry.

### Developability profiling

All 160 bispecifics and 71 monospecifics (65 parents, 6 additional controls) were profiled on the PROPHET-Ab high-throughput developability platform. All methods were followed as described in Arsiwala et al., 2025 unless specified otherwise.

*Affinity-capture self-interaction nanoparticle spectroscopy (AC-SINS)* measures self-interaction propensity as a plasmon wavelength shift (Δλmax) of antibody-coated gold nanoparticles relative to a reference (Sule et al., 2011; Liu et al., 2014). Measurements were collected in three buffer conditions: (i) phosphate-buffered saline (PBS), pH 7.4; (ii) 20 mM histidine, 150 mM arginine·HCl, 0.05% polysorbate 80, pH 6.0 (His/Arg formulation buffer); or (iii) 20 mM histidine, 150 mM NaCl, pH 6.0 (His/NaCl formulation buffer). Relative to the method published in Arsiwala 2025, we removed an intermediate centrifugation step which resulted in systematically elevated AC-SINS readouts for GDPa4 (current study) compared to GDPa1 (previous work).

*HPLC chromatography* comprises four methods: heparin affinity chromatography (HAC, retention time), hydrophobic interaction chromatography (HIC, normalized retention time), size-exclusion chromatography (SEC, percent monomer), and standup-monolayer affinity chromatography (SMAC, retention time).

*Intact mass spectrometry (IntactMS)* analysis used a rapid SEC-LC-MS workflow on a Thermo Fisher Vanquish Core HPLC coupled to a Thermo Fisher Orbitrap Exploris 240 mass spectrometer with the BioPharma option. Antibody samples were normalized to 0.25 mg/mL, and 2 µL was injected onto a Waters 125 Å, 2.5 µm, 4.6 × 30 mm desalting/guard column (Cat# 186011356). Separation was performed under isocratic conditions using 7% acetonitrile, 3% isopropanol, and 0.3% formic acid in water at 0.32 mL/min, with a total run time of 1.5 minutes per sample. Raw files were processed in BioPharma Finder, where spectra were deconvoluted for peak identification and reporting. Heterodimer fraction was then quantified from relative deconvoluted peak intensities against homodimer, half-antibody, and chain-mispaired species.

*Polyreactivity (PR)* was measured against three orthogonal reagents separately- CHO cell lysate (SMP:SCP::1:1), ovalbumin, and BVP. The final readout is reported as raw polyreactivity scores (pr_score). A normalized variant (pr_score_norm) was also measured but excluded from downstream analysis as near-collinear with the raw score. PR CHO and PR Ova methods have been previously described in Arsiwala et al., 2025. The baculovirus particle (BVP) ELISA was adapted from Hotzel et al., with SuperBlock (Thermo Fisher Scientific) substituted for BSA as the blocking buffer to avoid BSA-binding artifacts (**Figure S9**). BVPs (Medna Scientific, Inc., NC1999901) were coated at 0.25% (v/v) in 50 mM sodium carbonate buffer, pH 9.6. After blocking and PBS-Tween washes, test antibodies were added at 100 nM final concentration, incubated 1 h at room temperature, and detected with anti-human IgG-HRP followed by TMB substrate (reaction stopped with 2 M sulfuric acid). Absorbance at 450 nm was measured on a PHERAstar reader (BMG Labtech) and normalized to buffer-only blank wells; normalized values are reported as BVP scores.

*Thermostability (PTS-IF/nanoDSF)* monitors intrinsic tryptophan fluorescence ratio (F350/F330) during a thermal ramp to report three unfolding transitions: first melting temperature (Tm1), second melting temperature (Tm2), and onset of unfolding temperature (Tonset).

Atezolizumab, trastuzumab, and adalimumab served as end-to-end plate controls, run on every plate with approximately double the replicate count. Of these, only atezolizumab is a bispecific parent; trastuzumab and adalimumab were used solely for plate-level quality control and are not included in the 65-parent analysis set.

### Reproducibility across production campaigns validates the 65 parental antibodies

Combining data across production campaigns requires that the measurement platform produce consistent results. The 65 parent antibodies in this study were drawn from the published GDPa1 panel of 246 IgGs (Arsiwala et al., 2025). From these, 49 are the same isotype (IgG1) as used here. Both campaigns used similar PROPHET-Ab assay protocols. They differed in production date, cell line expansion, purification batch, manufacturing site. This overlap provides a direct test of reproducibility across campaigns for seven overlapping biophysical readouts (**Figure S1**). For HAC, SMAC, HIC, PR (Ova and CHO), and AC-SINS (PBS, pH 7.4 and His/Arg, pH 6), Pearson correlation coefficients ranged from 0.86 to 1.00 (n ≥47 for all but HAC [n = 23]).

The favorable correlations across campaigns for the overlapping seven developability properties reflect a combination of inherent measurement error and systematic process differences. They likely overestimate the measurement error within a single campaign. Accordingly, these results provide a useful baseline for distinguishing true variation in the properties of bispecific antibodies compared to the (monospecific) parental antibodies. All 160 bispecific and 71 (65 parents and 6 controls) monospecific antibodies reported here were evaluated together in a single campaign, further reducing random error in the comparisons described in this study.

### Data consolidation and processing

#### Data File

The supplemental data file contains:

1. "Definitions" sheet describing each column header corresponding to a data output;
2. "mAb Sequences" sheet documenting the amino acid sequences for all 71 IgGs produced in this work, the highest clinical status reached by the antibody, as of Feb 2025 as per <u>Thera-SAbDab</u> database, and the current approval status;
3. "bsAb Sequences" sheet documenting the amino acid for all 160 bispecific antibodies produced in this work. Antibody names follow the convention “A x B” for a bispecific with Fv from IgG A parent 1 and Fv from IgG B as parent 2;
4. "Assay Data -tidy format" sheet documents all data generated in this work in a tidy data format (one row per replicate) replication and conditions specified (e.g. buffer) where appropriate;
5. "Assay Data -median" sheet summarizing median, standard deviations and number of replicates for each developability assessment (one row per antibody);
6. “Production QC” sheet summarizing vendor production and purification QC. Included are details on yield, expression volume, purification stages, analytical SEC, nonreducing SDS-PAGE, and endotoxin levels for all antibodies.

#### Data processing

Per-antibody summary values were computed as the median across replicates. Median was chosen over mean because AC-SINS delta Lmax exhibits discrete plate-level outliers (single wells deviating by 4–10 nm from the remaining replicates of the same antibody) that contaminate arithmetic means. The same aggregation policy was applied uniformly across all assays.

### Developability Modeling

#### In silico features

Six computational predictor sources were applied to the monospecific parents. All six were computed on the GDPa1 panel and mapped to the corresponding GDPa4 monospecific antibodies (van Niekerk et al., 2026).

The six sources are: Aggrescan3D (aggregation propensity) (Kuriata et al., 2019), AntiFold (AlphaFold2-based structural features) (Hoie et al., 2025), DeepViscosity (viscosity and hydrodynamic radius prediction) (Kalejaye et al., 2025), MOE (molecular descriptors: molecular weight, isoelectric point, solvent-accessible surface area, hydrogen-bond counts) (Chemical Computing Group, 2024), SaProt_VH (VH domain embeddings via SaProtV2) (Su et al., 2024), and TAP (therapeutic antibody profiler) (Raybould et al., 2019).

Per-bispecific in silico features were derived by applying five compositional operators to each parent pair’s per-feature values: arithmetic mean, minimum, maximum, geometric mean (for positive-valued features only), and absolute difference (|parent_A -parent_B|, an order-invariant asymmetry measure).

#### Compositional baselines

For each of the 17 measured bispecific properties, we predicted the bispecific value from the two parent monospecific values using five operators: arithmetic mean, minimum, maximum, geometric mean, and absolute difference. Agreement between predicted and observed bispecific values was assessed by Spearman rho and R-squared (n = 158–160 per property).

#### Predictive modeling

Seven model families were evaluated: RidgeCV (alphas: 0.01, 0.1, 1, 10, 100), LassoCV, ElasticNetCV (L1 ratios: 0.1, 0.5, 0.7, 0.9, 0.95; same alpha grid), Random Forest (200 trees, max_features = sqrt(p), min_samples_leaf = 5), XGBoost (200 estimators, learning rate = 0.05, max depth = 4, subsample = 0.8, colsample_bytree = 0.8), HistGradientBoosting (200 iterations, learning rate = 0.05, max leaf nodes = 15), and partial least squares (PLS, n_components = min(p, 5)).

Six feature configurations defined the input space for each label:

1. Compositional baseline: a single feature, the arithmetic mean of the two parents’ values for the matching assay.
2. Corresponding experimental: all five compositional operators on the matching measured assay.
3. All experimental: all five operators across all measured assays.
4. In silico only: all five operators across all in silico sources.
5. In silico plus corresponding: configuration 4 combined with configuration 2.
6. In silico plus all experimental: configuration 4 combined with configuration 3.

In addition to these compositional operators, we also performed analysis when models were provided with the values for each parent individually (e.g. f(AC-SINS parent A, AC-SINS parent B)).

Linear models (Ridge, Lasso, ElasticNet, PLS) used a preprocessing pipeline of median imputation followed by standard scaling. Random Forest used median imputation without scaling. XGBoost and HistGradientBoosting handled missing values natively and received no imputation. All imputation was fitted per fold on training data only to prevent test-set leakage.

Labels with skewed distributions or hard boundaries received monotonic target transforms via scikit-learn’s TransformedTargetRegressor, which auto-inverts predictions so that all metrics are computed on the original measurement scale. The transforms were: SEC percent monomer received logit(y/100); BVP normalized score received logit(y); AC-SINS delta Lmax in His/Arg pH 6 and His/NaCl pH 6 received log(1 + y); SMAC retention time received log(y). All other labels used identity (no transform). Boundary values were clipped to (epsilon, bound -epsilon) with epsilon = 10^-6 to ensure finite transformed values.

#### Cross-validation strategy

Cross-validation enforced a parent-leakage guarantee: no test bispecific shares a parent with any training bispecific.

#### Parent-disjoint leave-one-out (LOO)

For each bispecific with parents A and B, the test set was that single bispecific; the training set comprised all bispecifics whose parents are neither A nor B. All 160 bispecifics received held-out predictions (subject to label observability). Typical training sets contained approximately 140 bispecifics.

Spearman rho (headline), Pearson r, R-squared, mean squared error (MSE), root mean squared error (RMSE), and mean absolute error (MAE) were computed on the original (untransformed) scale.

### Format-effect decomposition with repeated parents

For each assay, we computed residuals from the compositional mean baseline: r_ij = observed_ij -mean(parent_i, parent_j). We then fit an ordinary least squares (OLS) regression on a binary design matrix: r_ij = beta_i + beta_j + epsilon, where each bispecific row has indicator variables (0 or 1) for its two parent Fvs. The coefficient beta_i estimates the per-Fv format effect: the systematic deviation from the compositional mean attributable to that variable region when placed in bispecific context (**Figure S6**).

Per-Fv significance was assessed by t-test with Benjamini-Hochberg false discovery rate (FDR) correction. Only Fvs with degree of at least three (appearing in three or more bispecific combinations) were included; Fvs with fewer connections lack the statistical power for reliable coefficient estimation.

The model is fit without an intercept, so each beta_k has an absolute interpretation: beta_k = 0 means Fv k contributes no systematic format shift, and positive beta_k means bispecifics containing Fv k measure higher than the compositional mean predicts. An explicit intercept column cannot be added because it is a linear combination of the indicator columns (each row sums to exactly two). To test whether the bispecific scaffold introduces a global format bias that shifts all bispecifics in the same direction, we decomposed each coefficient as beta_k = mu + delta_k, where mu is the degree-weighted mean of all coefficients, and tested H0: mu = 0 using the delta method on the OLS covariance matrix.

### Statistical analysis and software

All analyses were implemented in Python 3.11 using scikit-learn (Pedregosa et al., 2011), XGBoost (Chen & Guestrin, 2016), and statsmodels (Seabold & Perktold, 2010). Figures were generated with matplotlib (Hunter, 2007) and seaborn (Waskom, 2021). Spearman ⍴ served as the headline ranking metric; Pearson r assessed linearity.

## Disclosure Statement

P.M.T. is a consultant for Ginkgo Bioworks, Inc. All other authors are past or present employees of Ginkgo Bioworks, Inc., who funded this work.

---

**Figure S1.**
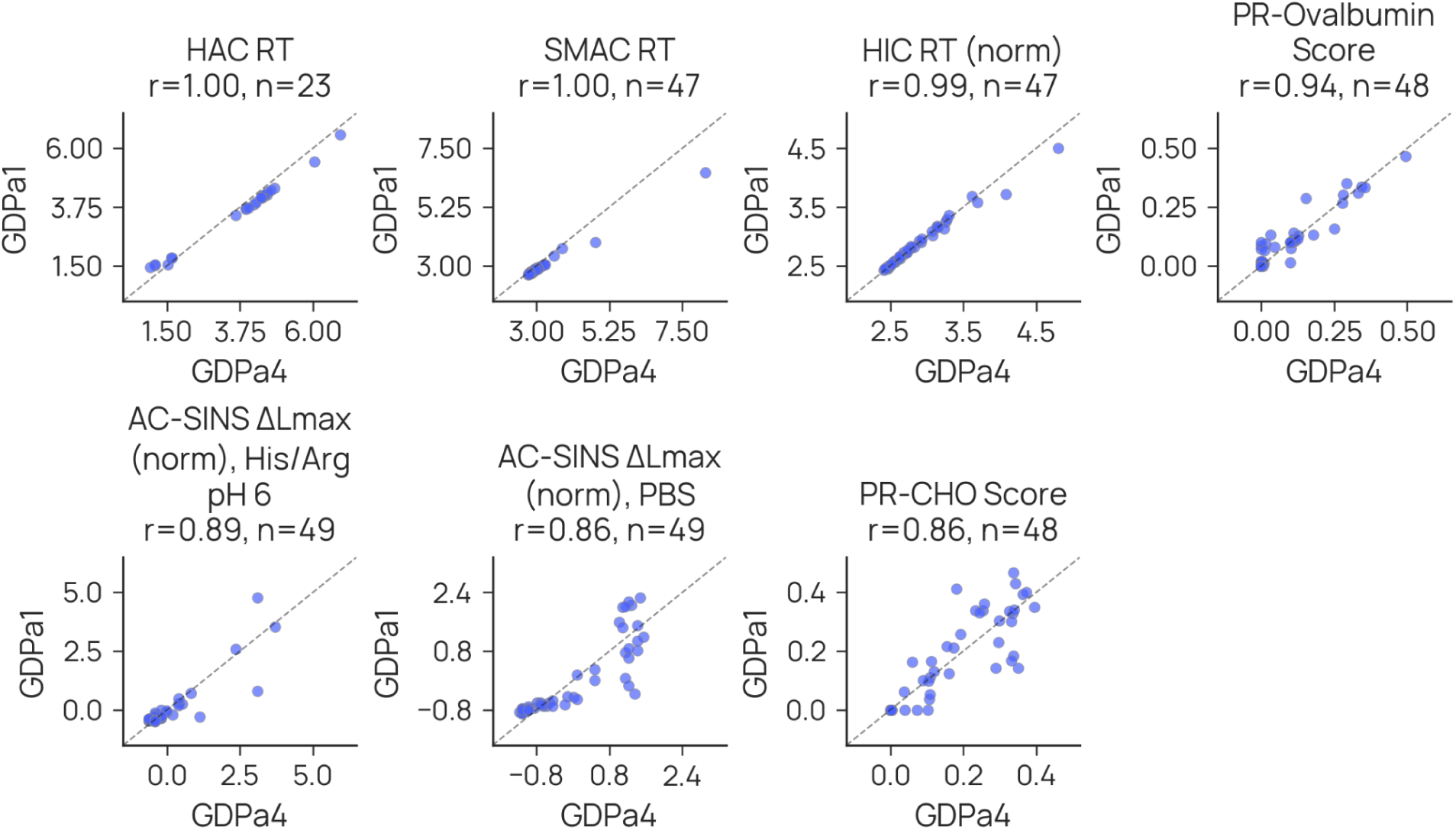
Reproducibility of antibody developability property profiling. Antibodies from this study (GDPa4; all reformatted as IgG1) are compared to the GDPa1 antibodies reported by Arsiwala et al. (2025). Of note, a method change was implemented for AC-SINS since the prior publication, resulting in elevated measures for this campaign.

**Figure S2.**
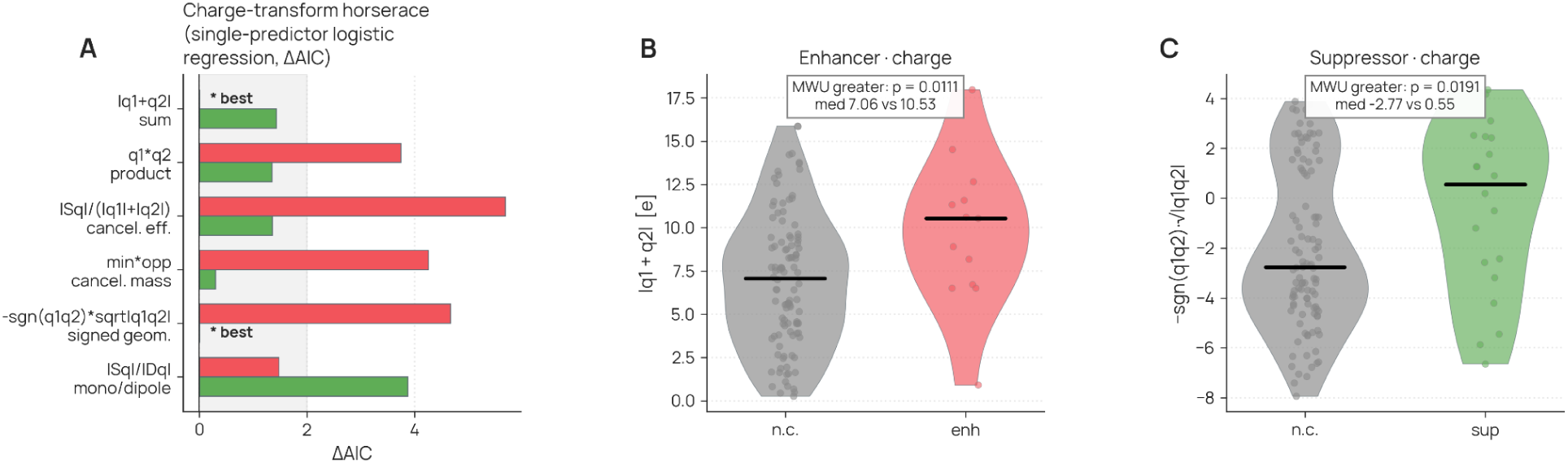
AC-SINS self-association in PBS pH 7.4. **(A)** Single-predictor logistic-regression horserace over charge transforms ranked by ΔAIC, with enhancer **(B)** and suppressor **(C)** charge distributions under their respective best transform.

**Figure S3.**
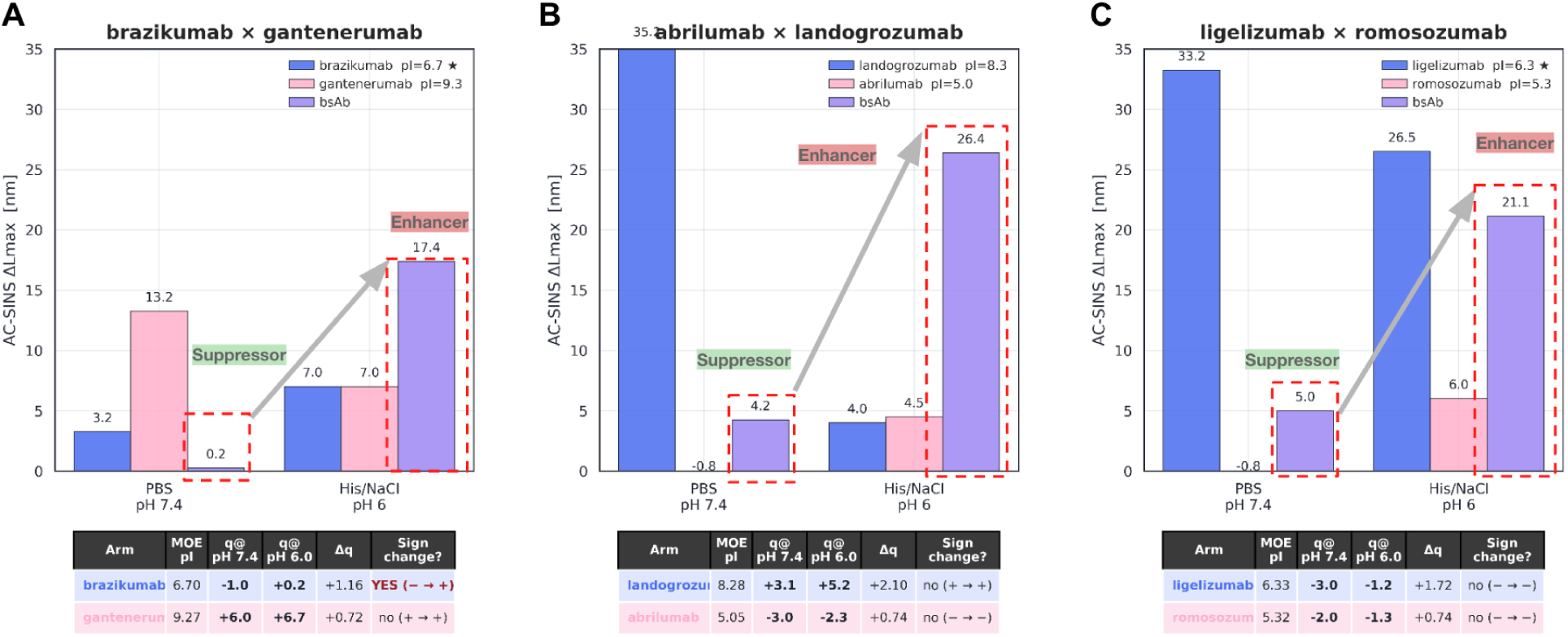
Emergent enhancer behavior by buffer context for AC-SINS. Expanding from analysis on a single buffer, emergence is presented in the context of 3 bispecific antibodies. **(A)** Bispecific antibody built of brazikumab and gantenerumab exhibits a large increase in AC-SINS signal when moving to pH6 in His/NaCl buffer, whereas the average for the constituent mAbs lowers. **(B)** Bispecific antibody built of abrilumab and landogrozumab exhibits large increase in AC-SINS signal despite dramatic decrease in average for each constituent mAb. **(C)** Bispecific antibody built of ligelizumab and romosozumab exhibits increase, potentially providing an example of the loss of suppressor behavior from the pair under context shift.

**Figure S4.**
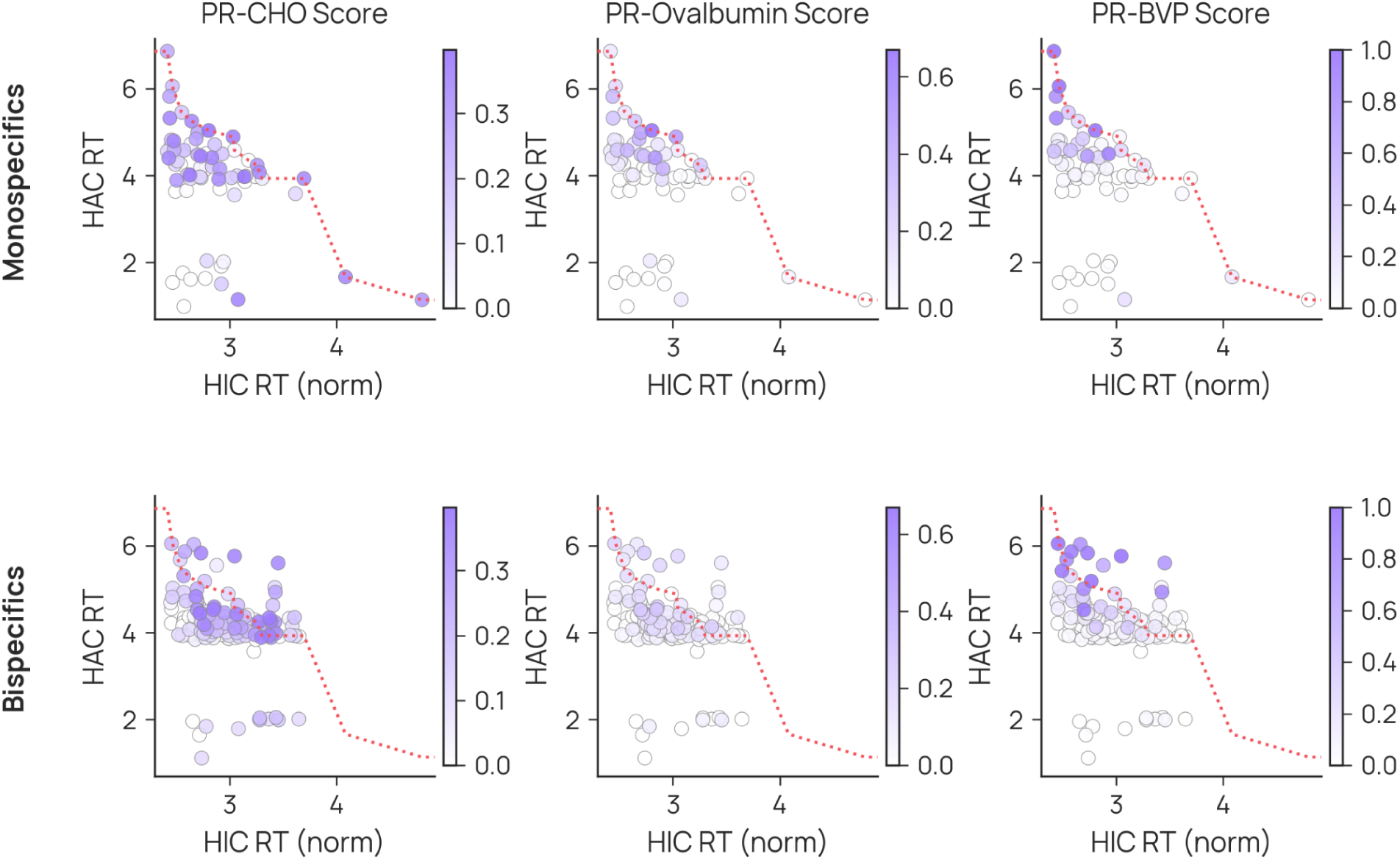
Class II polyreactivity over the HIC x HAC surface-property plane. (Top) The monospecific parents with a red dotted line highlighting the pareto frontier for high HIC x HAC. **(Bottom)** The bispecific library with parental frontier overlaid.

**Figure S5.**
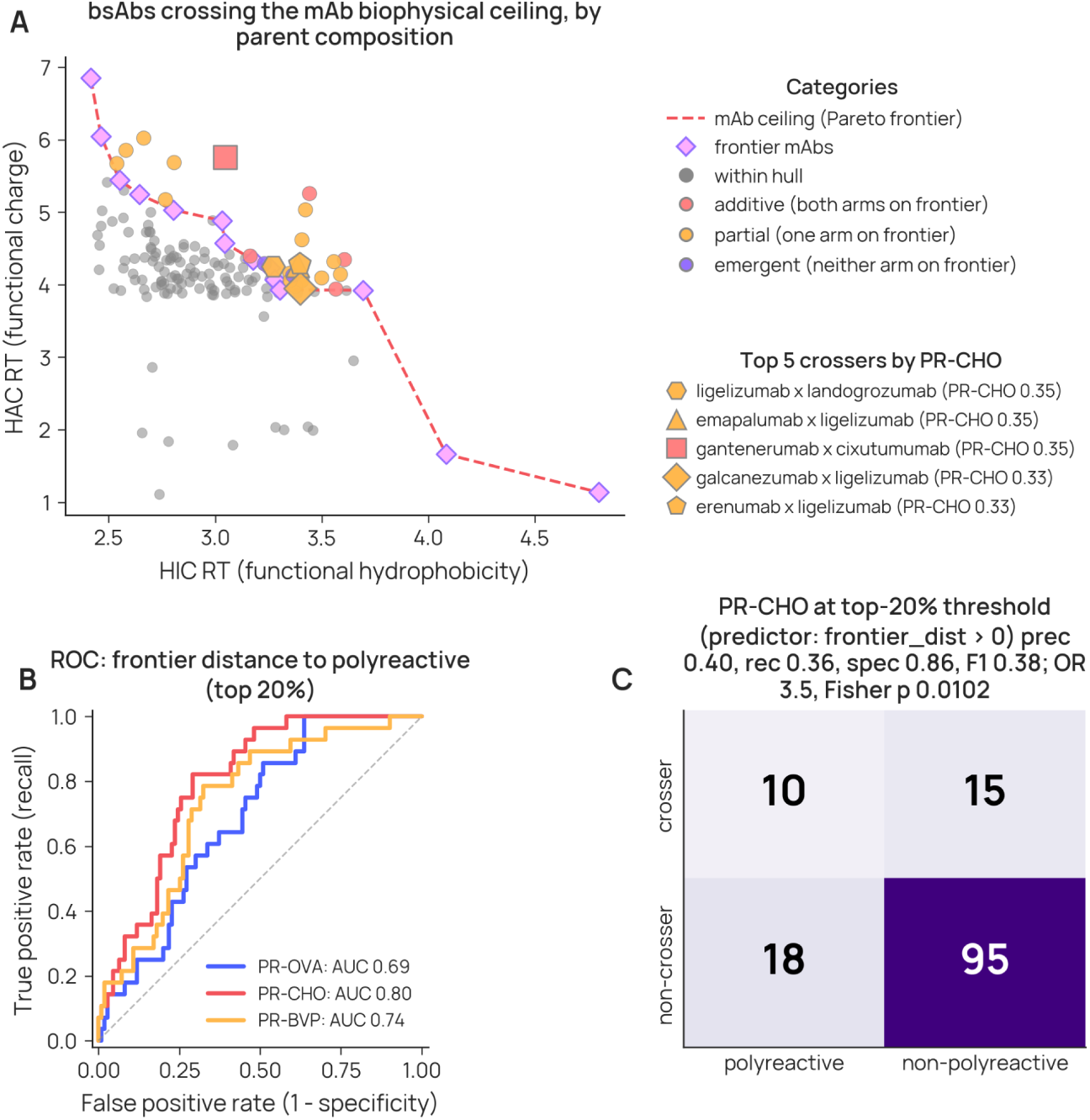
HIC x HAC frontier analysis. **(A)** Assessment of bispecific antibodies when mapped against the Pareto-frontier providing predictive assessment of PR-CHO. **(B)** Distance to this front, across polyreactivity assays, provides a measurable signal predictive of having elevated polyreactivity. **(C)** Bispecifics with parents that fall on the front and that exhibit crossing of the front are significantly more likely to be polyreactive.

**Figure S6.**
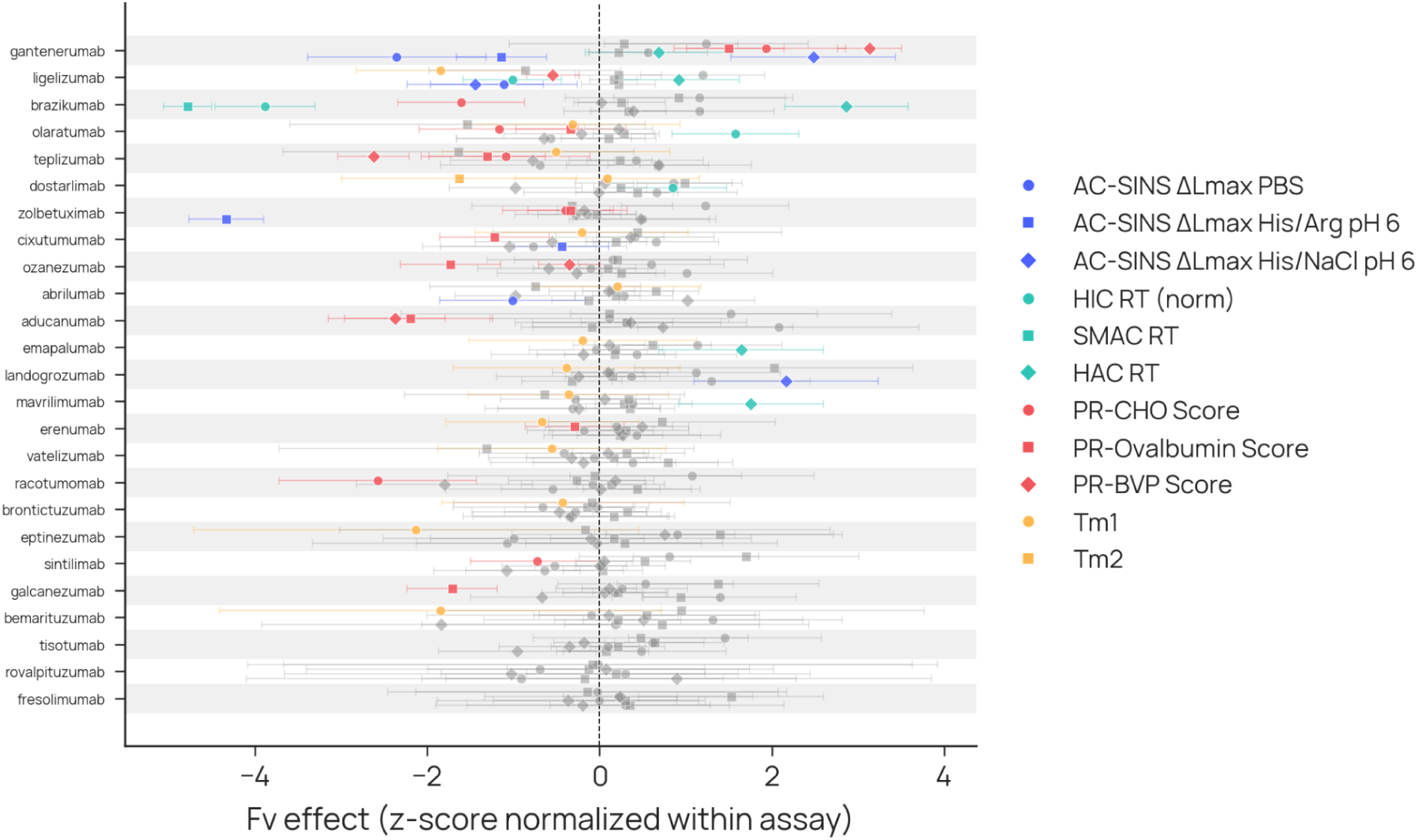
Per-Fv Format Effect Sizes. For those Fv present in at least 3 different bispecifics, a linear model of mean + Fv_a + Fv_b was fit. This network analysis permits the estimation of Fv effects for assays independent of the particular bispecific. The observation of Fv’s with effects statistically different than zero provide evidence for Fv’s which exhibit emergent behavior when no longer present as a two-Fv mAb.

**Figure S7.**
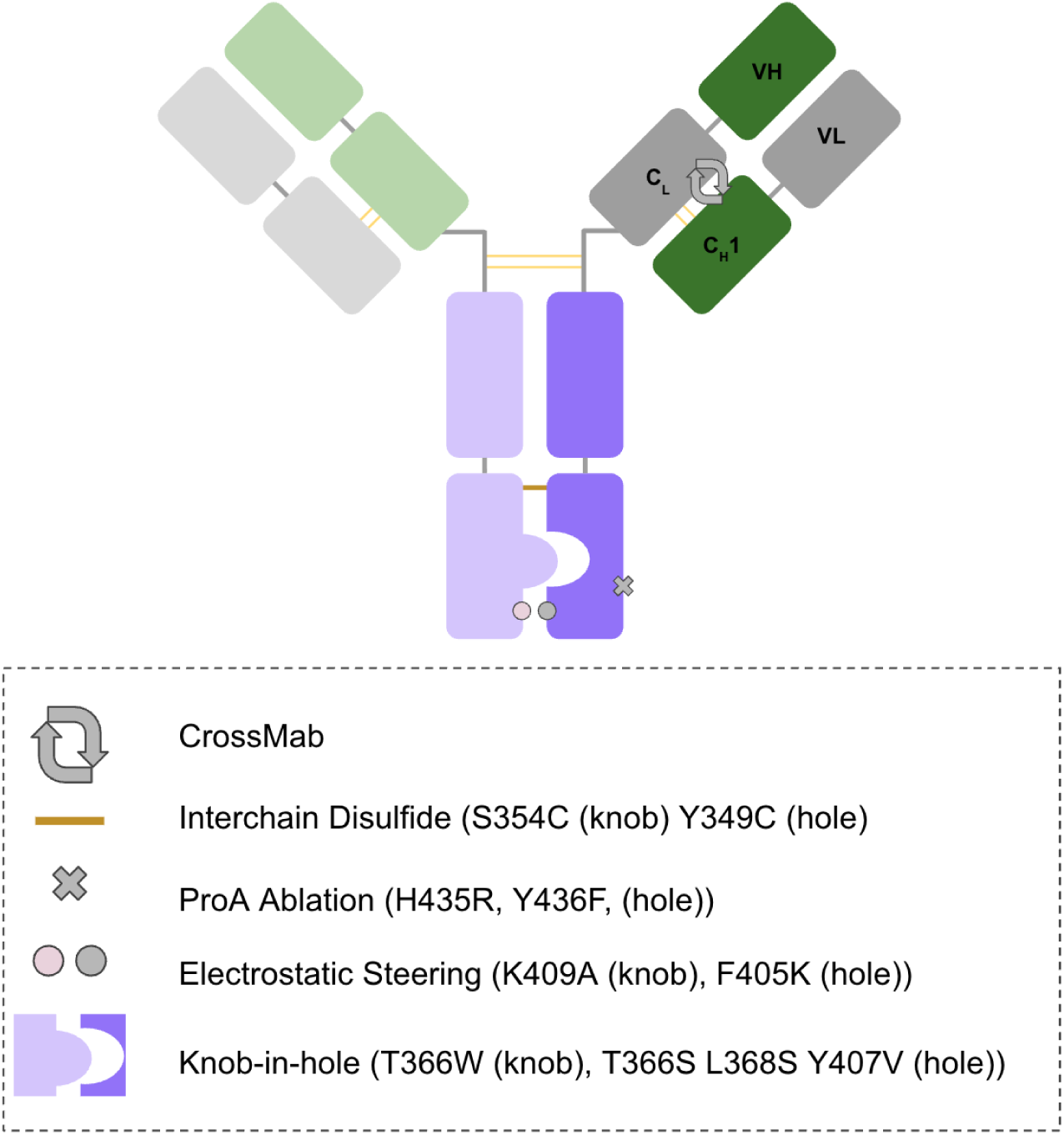
**Schematic of bispecific scaffold used in this study**

**Figure S8.**
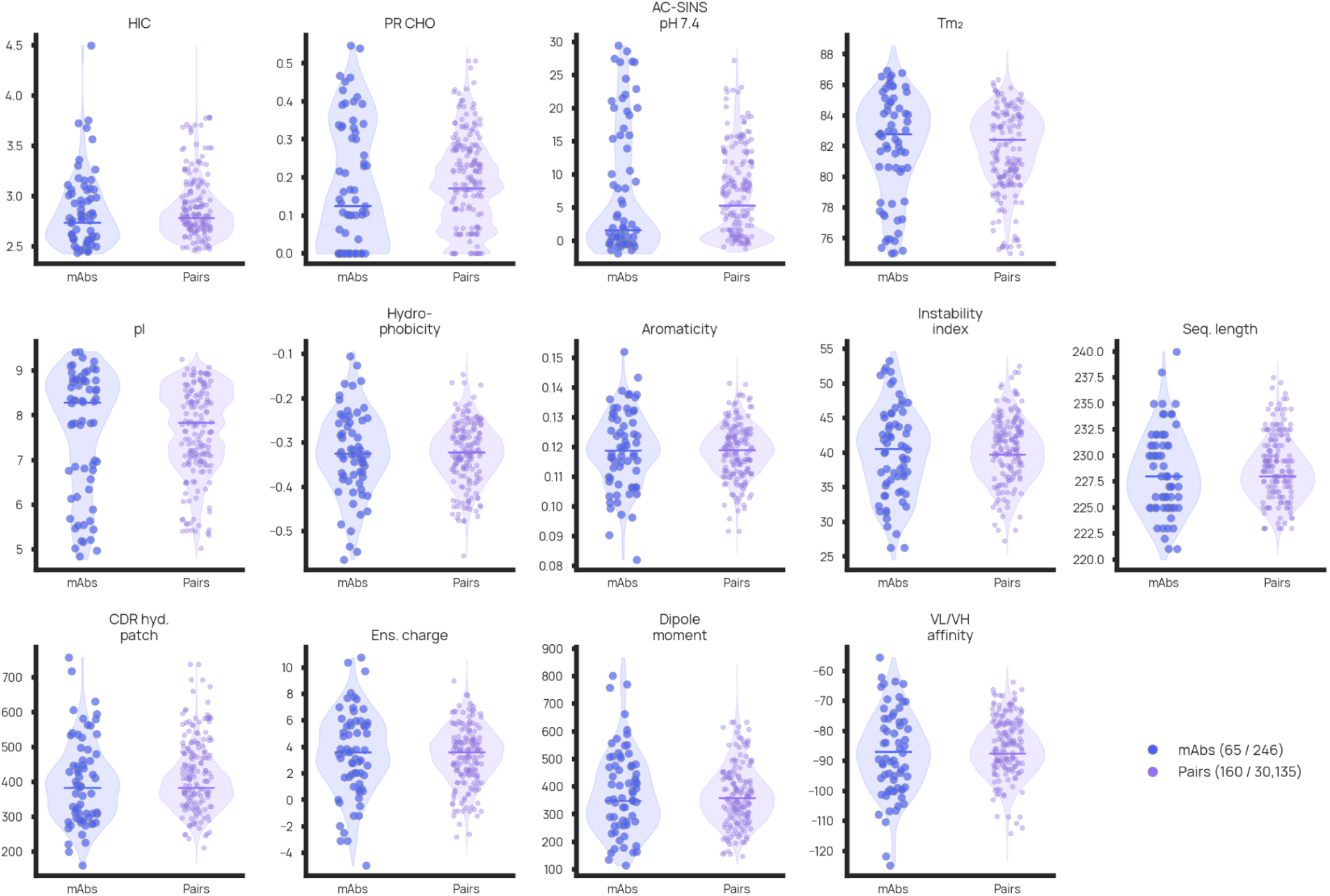
mAb property distributions. Assay readouts as provided in GDPa1, used in part as features for furthest point sampling. Each subplot shows two violins side by side: the full GDPa1 mAb population (blue, left) and all possible bispecific pair averages (purple, right), with the selected 65 parents and 160 pairs overlaid as scatter dots.

**Figure S9.**
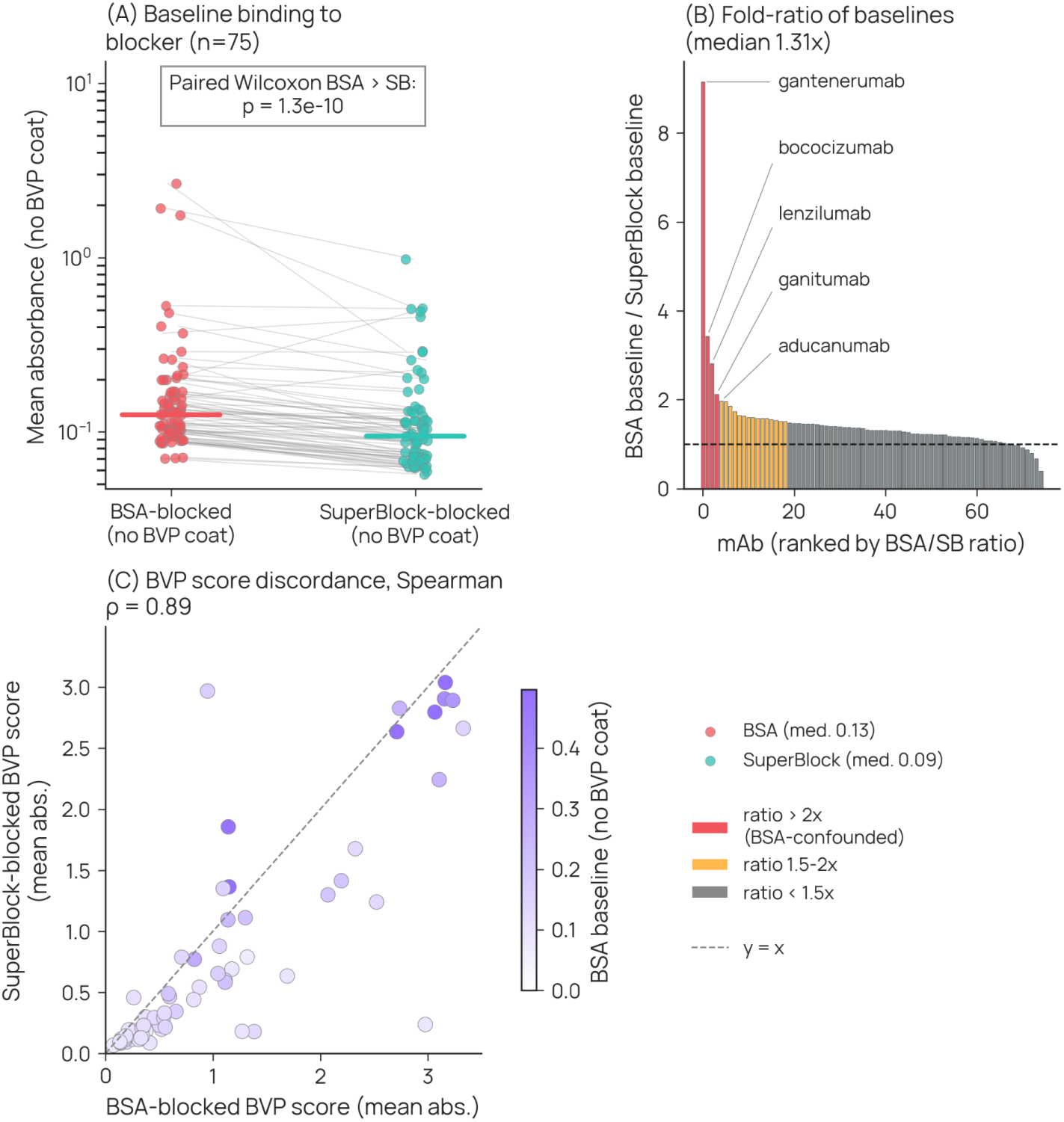
Influence of BSA and SuperBlock as blocking reagents in BVP ELISA assay. **(A)** Paired BVP assay results using either BSA (left) or SuperBlock (right) for blocking. **(B)** Ratio of baseline binding with labeling of mAbs exhibiting high BSA baseline compared with SuperBlock. **(C)** Correlation of BVP assay readouts with coloring by BSA baseline.

## Notes

https://datapoints.ginkgo.bio/dataset-access

### Funding

- Ginkgo Bioworks, Inc. (United States)

